# Requirement of a Wnt5A-Microbiota Axis in the Maintenance of Gut B Cell Repertoire and Protection from Infection

**DOI:** 10.1101/2024.03.08.584124

**Authors:** Soham Sengupta, Malini Sen

## Abstract

We investigated the influence of a Wnt5A-gut microbiota axis on gut B cell repertoire and protection from infection, having previously demonstrated that Wnt5A in association with gut commensals help shape gut T cell repertoire. Accordingly, Wnt5A heterozygous mice, which express less than wild type level of Wnt5A, and their isolated Peyer’s patches (PP), were studied in comparison with the wild type counterparts. The percentages of IgM and IgA expressing B cells were quite similar in the PP of both sets of mice. However, the PP of the Wnt5A heterozygous mice harbored significantly higher than wild type levels of microbiota bound B cell secreted IgA (sIgA), indicating the prevalence of a microbial population therein, that is significantly altered from that of wild type. Additionally, the percentage of PP IgG1 expressing B cells was appreciably depressed in the Wnt5A heterozygous mice in comparison to wild type. Wnt5A heterozygous mice, furthermore, exhibited notably higher than the wild type levels of morbidity and mortality following infection with *Salmonella enterica*, a common gut pathogen. Difference in morbidity/mortality correlated with considerable disparity between the PP-B cell repertoires of the *Salmonella* infected Wnt5A heterozygous and wild type mice, the percentage of IgG1 expressing B1b cells in the PP of heterozygous mice remaining significantly low as compared to wild type. Overall, these results suggest that a gut Wnt5A-microbiota axis is intrinsically associated with the maintenance of gut B cell repertoire and protection from infection.

**Importance:** Although it is well accepted that B cells and microbiota are required for protection from infection and preservation of gut health, a lot remains unknown about how the optimum B cell repertoire and microbiota are maintained in the gut. The importance of this study lies in the fact that it unveils a potential role of a growth factor termed Wnt5A in the safeguarding of the gut B cell population and microbiota, thereby protecting the gut from the deleterious effect of infections by common pathogens. Documentation of the involvement of a Wnt5A-microbiota axis in the shaping of a protective gut B cell repertoire, furthermore, opens up new avenues of investigations for understanding gut disorders related to microbial dysbiosis and B cell homeostasis that till date, are considered incurable.

## Main Text

### Observation

Gut microbiota and the immune network remain closely linked in the fight of the human host against infection by common pathogens (1, 2). Yet, the association of gut microbiota with immune cells in the context of immune defense remains unexplained. We previously demonstrated that Wnt5A in connection with gut commensals helps shape the gut immune T cell repertoire at the steady state (3). In view of the functional interdependence of B and T cells (4–6), we presently explored the possibility of an association of a Wnt5A-gut commensal axis with the shaping of gut B cell repertoire and thereby protection from infection. This investigation was further prompted by experimental findings indicating gut commensals as a potential regulator of antibody class switching, a major aspect in the development and maintenance of B cell repertoire (7–9).

For our study model we compared wild type mice with Wnt5A heterozygous mice (harboring one functional copy of the Wnt5A gene), where reduced Wnt5A expression with reference to wild type, correlates with altered abundance of the gut microbiota and a potential for dysbiosis (3). In both Wnt5A heterozygous and wild type mice, we quantified IgM/IgA/IgG1 expressing B cell subtypes (B220+: bone marrow derived mature B cells, and B1a/B1b: independent B cell lineage important for innate immunity) in the respective Peyer’s patches (PP). Both IgM and IgA expressing B cells were targeted due to the abundance of IgM and IgA in the gut (10–12). IgG1 expressing B cells were targeted on account of reports demonstrating IgG1 mediated immune protection of the gut against infection by *Salmonella*, a common gut pathogen (8, 13). Alongside, we measured bacteria bound IgA in the PP of all sets of mice in line with its strong correlation with alteration of microbial abundance and dysbiosis, as well as prevalence of gut infection (4, 14–19). Experiments performed in the steady state revealed significantly higher level of bacteria bound secreted IgA (sIgA) but similar percentages of IgM and IgA expressing B220+, B1a and B1b cells in the PP of the Wnt5A heterozygous mice as compared to the wild type counterparts (Figures 1A, 1B and 1C). The high bacteria bound sIgA level in the Wnt5A heterozygous mice validated alteration in gut bacterial abundance therein as compared to the wild type, in line with previously published results. (3, 15, 17–19). In view of the already documented increased association of potential pathogens such as *Helicobacter sp*. and *Prevotella sp*. with the PP of Wnt5A heterozygous mice as compared to the wild type (3), and the propensity of sIgA binding to potential pathogens (4, 15, 17–19), the current observation suggested higher probability of gut infection in the Wnt5A heterozygous mice as opposed to the wild type. Additional documentation of significantly depressed percentage of PP IgG1 expressing B cells (B1a, B1b and B220) in the Wnt5A heterozygous mice in comparison to wild type (Figure 1D) in the light of already proven IgG1 dependent protection against *Salmonella* infection (8, 13) furthermore, strengthened the possibility of increased infection of the Wnt5A heterozygous mice by *Salmonella*. The observed difference between Wnt5A heterozygous and wild type mice in B cell IgG1 expression could be due to variation in B cell class switching caused by altered abundance in gut microbiota. All gating of FACS pertaining to Figure 1 is demonstrated in Supplementary Figure 1.

**Figure 1:**
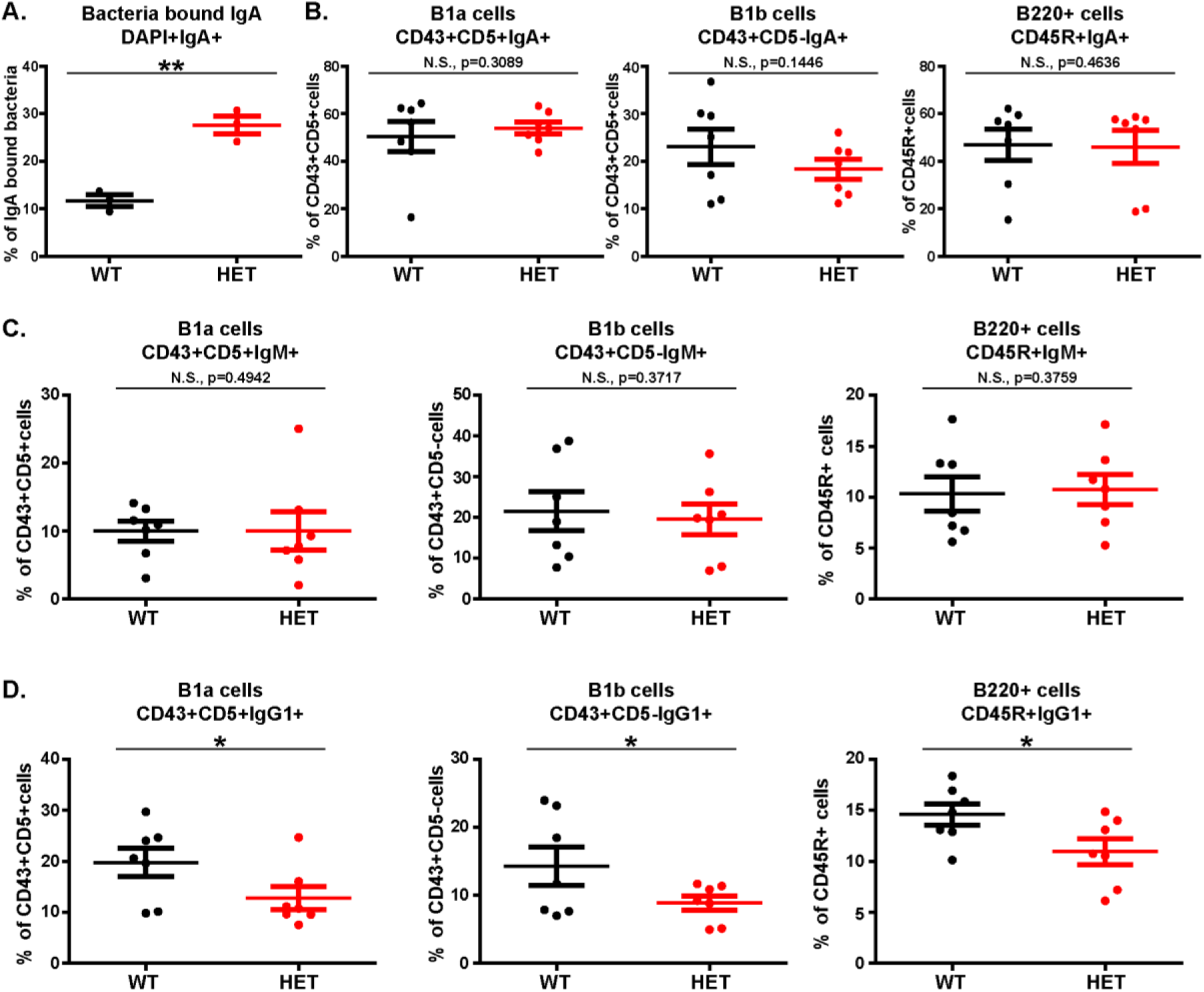
Wnt5A dosage controls the steady state level of gut (PP) sIgA bound bacteria and B cell repertoire. (A) Graph representing increased percentage of bacteria bound sIgA in the PP of Wnt5A heterozygous (HET) mice in comparison to wild type (WT) (n=3). (B & C) There is no significant change in the percentage of IgA expressing (B) and IgM expressing (C) B1a (CD43+CD5+), B1b (CD43+CD5-) and CD45R/B220+ B cells between the Wnt5A heterozygous and wild type mice, as represented graphically (n=7). (D) Plots showing significantly lower percentage of IgG1 expressing B1a, B1b and B220+ B cells in the PP of Wnt5A heterozygous mice as compared to wild type (n=7). Data represented as mean ± SEM. p≤ 0.05 was considered as significant statistically. Significance is represented by * in the following manner: * p≤ 0.05, ** p≤ 0.01, *** p≤ 0.001. “N.S.” denotes non-significant. “n” represents number of mice in each experimental set.

The likelihood of an increased susceptibility for gut infection in the Wnt5A heterozygous mice as opposed to the wild type led us to study and compare the outcome of *Salmonella enterica* infection in the Wnt5A heterozygous and wild type mice. Both sets of mice were infected to the same extent through oral gavage (10^7^ *Salmonella enterica* per mouse). In line with depressed IgG1 expressing B cells in the Wnt5A heterozygous mice in the steady state, we observed significantly higher infection induced morbidity (as noted by slower movement, lack of hunger and thirst) and mortality in the Wnt5A heterozygous mice as compared to the wild type mice, leading to lesser numbers of healthy mice in the heterozygous cohort as oppose to the wild type.(Figure 2A, 2B and 2C). High morbidity/mortality in the Wnt5A heterozygous mice was associated with higher than wild type percentages of PP associated IgA/IgM/IgG1 expressing B cell subtypes (higher IgA and IgM: B1a, B1b and B220; higher IgG1: B1a, B220) (Figure 2D, 2E, 2F), which could be responsible for inflammation without resolution of the deleterious effects of infection. On the other hand, the percentage of IgG1 expressing B1b cells, which are important for warding off bacterial infections along with B1a cells (20–25) remained much lower in the Wnt5A heterozygous mice than in the wild type (Figure 2F). The extent of sIgA bound bacteria in both wild type and Wnt5A heterozygous mice appeared the same (Figure 2G), suggesting that the seemingly increased B cell activation in the Wnt5A heterozygous mice in terms of IgA expression perhaps did not result in any increase in suppression of infection through bacterial binding of sIgA. It is quite possible that the observed morbidity/mortality in the infected wnt5A heterozygous mice is due to the continued prevalence of pathogenic conditions associated with infection and inflammation. All gating of FACS pertaining to Figure 2 is explained in Supplementary Figure 2.

**Figure 2:**
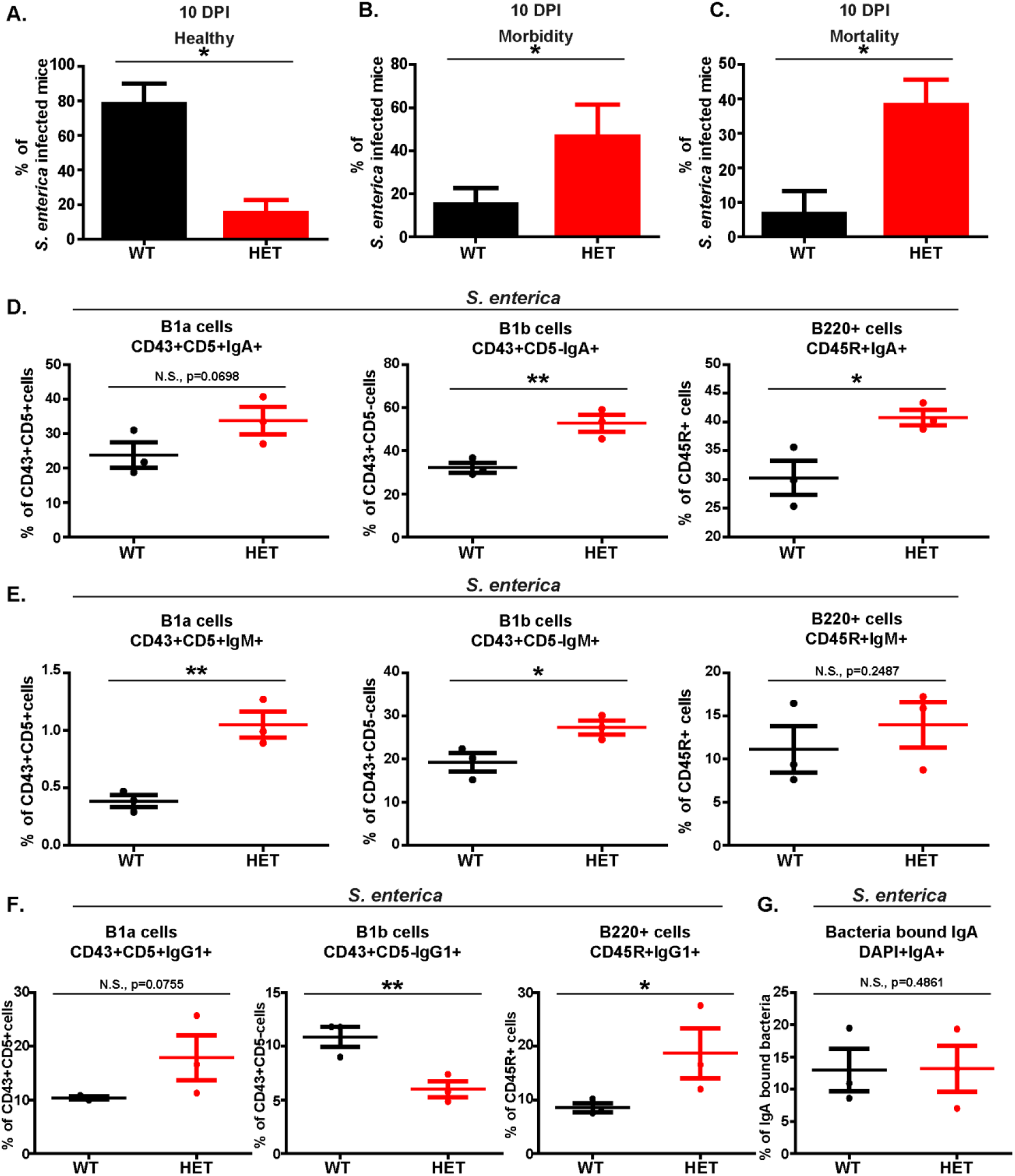
Wnt5A dosage dependent morbidity/mortality after *Salmonella* infection correlates with an altered B cell repertoire. (A, B & C) Graphs showing depletion in percentage of healthy Wnt5A heterozygous (HET) mice as compared to the wild type (WT) controls (A), due to significant increase in morbidity (B) and mortality (C) under *Salmonella enterica* infected conditions at DPI (Days Post Infection) = 10 (n=3 separate experiments with 4-5 mice in each WT/HET group). Mice demonstrating morbidity had slower movement, with lack of hunger and thirst. Mortality represents the percentage of mice that succumbed to death from the infection. (D & E) Graphs depicting higher percentage of IgA (D) and IgM (E) expressing CD45R/B220+, B1a (CD43+CD5+) and B1b (CD43+CD5-) cells in the PP of *Salmonella enterica* infected Wnt5A heterozygous mice in comparison to the wild type counterparts at 10 DPI. The increase in B1a cell IgA and B220 cell IgM however was not statistically significant (n=3). (F) Graphs demonstrating that despite increased percentages of IgG1-B220+ and IgG1-B1a (although not statistically significant) populations in the *S*.*enterica* infected PP of the Wnt5A heterozygous mice as compared to the wild type counterparts, *S. enterica* infected Wnt5A heterozygous mice PP had significantly lower percentage of IgG1-B1b cells as opposed to the wild type controls at 10 DPI (n=3). (G) Plot depicting no difference between sIgA bound bacteria in the PP of *S. enterica* infected wild type and heterozygous mice. Data represented as mean ± SEM. p≤ 0.05 was considered as significant statistically. Significance was represented by * in the following manner: * p≤ 0.05, ** p≤ 0.01, *** p≤ 0.001. “N.S.” denotes non-significant. “n” represents number of mice in each experimental set.

Overall these results indicate that the gut microbiota abundance/profile that is dependent on the dosage of Wnt5A is associated with the shaping of gut B cell repertoire. Alterations in the gut microbial population and associated B cell repertoire due to compromised levels of gut Wnt5A may lead to uncontrolled inflammation by pathogens, as often is the case during microbial dysbiosis. The potential role of the Wnt5A-gut microbiota axis in the maintenance of gut B cell repertoire and protection from the harmful effects of infection, as projected through this observation, opens up new avenues for further investigations into the role of Wnt5A signaling in gut health and disease.

## Materials and Methods

### Reagents

All reagents used in this study are enlisted in Table 1.

**Table 1:** All reagents used in this study are enlisted below.

| Reagent or Resource | Source | Ref. No. |
| --- | --- | --- |
| RPMI 1640 | Life Technologies (Thermofisher Scientific, USA) | 31800-022 |
| HI FBS | Life Technologies (Thermofisher Scientific, USA) | 10082147 |
| L-glutamine | Life Technologies (Thermofisher Scientific, USA) | 25030081 |
| DAPI | Life Technologies (Thermofisher Scientific, USA) | D1306 |
| DNase I | Life Technologies (Thermofisher Scientific, USA) | AM2222 |
| Collagenase D | Sigma-Aldrich, USA | 11088866001 |
| NaCl | Sigma-Aldrich, USA | S5886 |
| Fixable Viability Stain 450 | BD Biosciences, USA | 562247 |
| APC rat anti-mouse CD5 | BD Biosciences, USA | 550035 |
| PeCy7 rat anti-mouse CD43 | BD Biosciences, USA | 562866 |
| PeCy7 rat anti-mouse<br>CD45R/B220 | BD Biosciences, USA | 561881 |
| PerCP Cy5.5 rat anti-mouse<br>IgM | BD Biosciences, USA | 550881 |
| BV605 rat anti-mouse IgG1 | BD Biosciences, USA | 563285 |
| FITC rat anti-mouse IgA | BD Biosciences, USA | 559354 |

### Mice maintenance, PP isolation and infection

Wnt5A wild type (+/+) and heterozygous (+/-) mice belonging to the strain B6;129S7-*Wnt5a*^*tm1Amc*^*/*J obtained from Jackson Laboratory were bred and maintained in the institutional animal house facility of CSIR-IICB as previously described (3). Mice, 8 – 10 week old, used for control and experimental sets had similar male:female ratio. Cells from PP (Peyer’s patches) were isolated following previously published protocol (3). Briefly, the PP identified from the small intestine, were isolated as 1mm sections, shaken for 20 min at 37° C with 200 RPM in RPMI 1640 media containing 10% FBS and 10mM EDTA, following which media was changed with RPMI 1640 (+10% FBS) containing Collagenase D and DNase I for further 15 min shaking under similar conditions. After repeated dispersion by passage through 40 μm cell strainer and short-term shakings the almost dispersed single cells were used for experiments. Mice were infected with *Salmonella enterica* purchased from MTCC, Chandigarh, India (MTCC 3224) at dosage of 10^7^ bacteria per mice via oral gavage.

### Flow Cytometry

Cells were prepared for flow cytometry following previously published protocols (3). Briefly, PP-cells were incubated in 1% BSA (blocking agent) and then incubated at 4° C with fluorophore conjugated antibodies dissolved in 0.5% BSA in PBS for 1 hr. B cell subset identification was done with appropriate antibodies using flow cytometry. For studying bacteria bound sIgA, PP cells were maintained overnight in antibiotic free RPMI 1640 medium supplemented with 10% FBS and 2mM L-glutamine. The harvested sup was first centrifuged for 5 min at 700 g to remove floating eukaryotic cells, and centrifuged again at 7155g for 3 min to pellet down bacterial cells. The pellet was suspended in 0.5% BSA and PBS and stained with fluorophore bound anti-IgA antibody, following counterstaining with DAPI (bacteria identification). Data acquisition was done using BD.LSR Fortessa Cell analyzer and flow cytometric data analysis was done by FCS Express 5 software. All cell percentages are calculated and presented on the basis of the previous gate.

### Statistical analysis

Statistical analysis was done utilizing Paired or Unpaired Student t test as per need using Graphpad Prism 5 software. Graphs and line diagrams are represented as mean ± SEM and p≤ 0.05 was considered statistically significant. Significance was represented by * in the following manner: * p≤ 0.05, ** p≤ 0.01, *** p≤ 0.001.

## Ethics Statement

All animal experiments were conducted with approval of Animal Ethics committee of CSIR-IICB as per meeting IICB/AEC/Meeting/Sep/2019/1 held on 19/09/2019.

## Acknowledgments

The authors acknowledge Payel Karmakar Halder for FACS, CSIR-IICB Instrument Facility for instrument support and CSIR-IICB animal house facility for animal breeding/maintenance.

The authors acknowledge financial support from Institutional funding. SS is supported by fellowship from CSIR, Government of India. Funding agencies played no role in designing or execution of the study or in decision of manuscript submission. The authors declare no Conflict of Interest related to this study.

